# Untrained Convolutional Neural Networks as Feature Extractors for Structural MRI

**DOI:** 10.64898/2026.06.07.730652

**Authors:** Arel Encin, Inés Gonzalez Pepe, Yohan Chatelain, Erin Dickie, Tristan Glatard

**Affiliations:** Krembil Centre for Neuroinformatics, Centre for Addiction and Mental Health, Toronto, ON M5T 1R8, Canada; University of Toronto, Toronto, ON M5S 1A1, Canada; Department of Psychiatry, University of Toronto, Toronto, ON M5S 1A1, Canada; Institute of Medical Science, University of Toronto, Toronto, ON M5S 1A1, Canada; Department of Computer Science, University of Toronto, Toronto, ON M5S 1A1, Canada; Department of Computer Science and Software Engineering, Concordia University, Montreal, QC H3G 1M8, Canada

**Keywords:** Untrained Convolutional Neural Networks, structural MRI

## Abstract

We demonstrate that features extracted from structural MRI using un-CNN, an untrained convolutional neural network, achieve predictive performance comparable to or exceeding that of state-of-the-art pretrained foundation models across three structural MRI datasets and three downstream tasks. Un-CNN extends a classical 3D CNN architecture with multi-channel inputs, a hierarchical encoder with multi-scale feature aggregation, and covariance pooling. Untrained CNNs circumvent several key limitations of trained models, including high computational cost and memory requirements, the need to distribute large model weights, risks of data leakage, and challenges in reproducibility.

## Introduction

Extracting features from brain structural Magnetic Resonance Images (sMRI) is a fundamental requirement of predictive modeling. Historically, feature extraction has relied on handcrafted morphometric pipelines such as FreeSurfer (1), which perform cortical surface reconstruction and parcellation to produce predefined anatomical measurements including cortical thicknesses, surface areas, and regional volumes. While biologically interpretable, these features require hours of computation per scan and lengthy quality control.

Deep learning (DL) approaches subsequently implemented feature extraction by training convolutional neural networks (CNNs) and other model architectures directly on MRI volumes, learning task-specific representations through supervised learning (2, 3). More recently, foundation models have extended this paradigm by pretraining model on large MRI datasets (4,000 to 75,000 scans) using self-supervised learning, producing features transferable to downstream tasks (4–7). However, all three paradigms share substantial computational burdens: handcrafted pipelines require hours per scan, DL models demand intensive training, and foundation models additionally necessitate distributing pretrained weights in the order of gigabytes. Retraining or fine-tuning these models when architectures change or new data becomes available remains cumbersome and resource-intensive.

While benchmarking sMRI foundation models in (8), we serendipitously observed that an untrained CNN with randomly initialized weights achieved performance competitive with FreeSurfer on brain age prediction and sex classification. This finding connects to prior work demonstrating the utility of untrained DL networks in other domains: in 2011, Saxe et al. (9) showed that untrained CNNs can achieve surprisingly competitive performance on object recognition tasks, suggesting that the architectural structure of CNNs itself imposes useful inductive biases. More recently, untrained models have shown promise in MRI reconstruction (10), and activation in untrained CNNs was showed to align strongly with activation in the visual cortex (11). However, no prior work has explored untrained CNNs as feature extractors for structural brain MRI phenotypic prediction.

Beyond predictive performance, untrained CNN feature extraction offers several practical advantages over existing approaches. Computation is substantially faster, requiring only a single forward pass and no training loop. Model sharing is trivial as the entire model is defined by code and a random seed, eliminating the need to distribute large model files. Memory requirements are modest since no training state, gradient buffers, or optimizer parameters are stored. Computational reproducibility is facilitated as the same seed generally produces identical features across execution environments. Finally, untrained models carry no risk of data leakage, as weights have never been exposed to any brain data, a concern that is increasingly relevant as foundation models are pretrained on datasets that may overlap with evaluation cohorts.

This paper makes the following contributions: (1) we propose *un-CNN*, a 3D untrained CNN architecture incorporating multi-scale readout, depthwise separable convolutions, multi-channel preprocessing, and covariance pooling specifically designed for random-weight feature extraction from brain structural MRIs; (2) we evaluate this architecture against four state-of-the-art brain MRI foundation models (AnatCL (4), 3D-Neuro-SimCLR (5), BrainIAC (6), SwinBrain (7)) and FreeSurfer baselines, and across three large publicly available datasets: the Nathan Kline Institute–Rockland Sample (NKI) (12), the Healthy Brain Network (HBN) (13), and the Parkinson’s Progression Markers Initiative (PPMI) (14). These datasets span children, adolescents, adults, and elderly clinical populations. We evaluated the models using three downstream tasks: sex classification, brain age regression, and body-mass index prediction. Our untrained CNN models are pub-licly available at https://github.com/epics-lab/untrained-cnn and https://huggingface.co/epics-lab/untrained-cnn.

### Untrained Convolutional Neural Network Architecture

The un-CNN architecture, illustrated in Figure 1, was developed from the architecture presented in (15) for CT scan classification, which consisted of four blocks, each containing a convolutional layer, a max-pooling layer, and a batch normalization layer, with 64, 64, 128, and 256 filters, respectively. The original architecture included a global average pooling layer followed by a fully connected layer, producing a single 512-dimensional image representation from the final block only. We extended this initial CNN architecture into a four-block 3D CNN with three components: multi-channel input construction, a hierarchical encoder, and multi-scale readout with covariance pooling. In contrast to the original single-block readout, our architecture extracts features from every block, capturing both mean activations and pairwise channel covariances, and concatenates them into a 9,664-dimensional embedding.

**Fig. 1.**
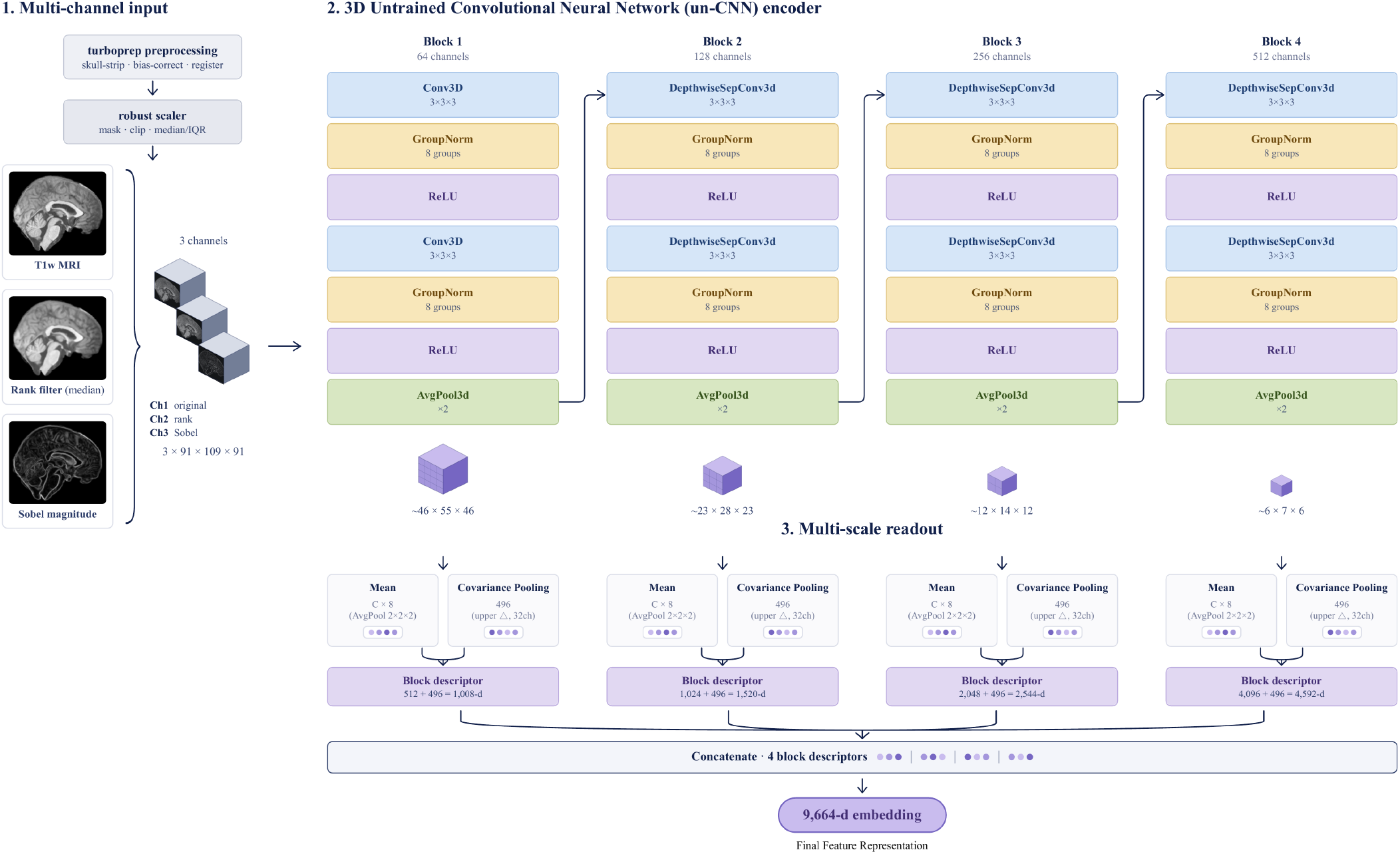
Overview of the proposed untrained 3D CNN (un-CNN) feature extraction pipeline. (1) Multi-channel input: each TurboPrep-processed volume undergoes robust intensity scaling, then three channel variants (original, rank-filtered, Sobel magnitude) are stacked into a 3 *×* 91 *×* 109 *×* 91 input tensor. (2) Hierarchical encoder: four blocks with randomly initialized, frozen weights progressively increase channel depth (64, 128, 256, 512) while halving spatial resolution. (3) At each block output, mean features (AvgPool to 2 *×* 2 *×* 2) and covariance features (upper triangle, 32 channels) are extracted and concatenated into a block descriptor. The four descriptors are concatenated into a 9,664-dimensional embedding.

#### A. Pre-processing

Pre-processing consists of the TurboPrep pipeline (16), which performs N4 bias field correction with the ANTs tool (17), skull stripping with SynthStrip (18), rigid registration to the MNI152 ICBM 2009c nonlinear symmetric template (193*×* 229 *×*193 voxels, 1 mm isotropic) (19, 20) with ANTs (21), resampling to 1 mm isotropic resolution also with ANTs, and intensity normalization with WhiteStripe (22), requiring approximately 1–2 minutes per scan. For the un-CNN, volumes were interpolated from the template resolution of 193*×*229 *×*193 to 91 *×*109 *×* 91 voxels. For foundation models requiring different input dimensions, volumes were interpolated to their respective model-specific resolutions.

#### B. Multi-channel input construction

After pre-processing and before the 3D image is passed to the model, each brain volume undergoes robust intensity normalization. Background voxels below the 5th intensity percentile are masked, while intensities within the brain mask are clipped at the 2nd and 98th percentiles, centered on the median, and scaled by the interquartile range. Three versions of the normalized volume are then generated: (1) the original intensity image, (2) a median rank-filtered image providing edge-preserving smoothing, and (3) a Sobel gradient-magnitude image emphasizing boundaries. These three volumes are stacked along the channel dimension to form a multi-channel input tensor of size 3*×* 91*×* 109*×* 91. The first `Conv3d` layer receives all three channels simultaneously and maps them to 64 output feature maps in a single convolutional operation.

#### C. Hierarchical encoder

The multi-channel input is processed through four sequential blocks with increasing channel depths of 64, 128, 256, and 512. Block 1 employs standard convolutions, while blocks 2–4 employ depthwise separable convolutions (23), in which each input channel receives an independent spatial filter prior to pointwise convolution 1*×* 1 that mixes information across channels. Each block applies two successive convolutional layers, followed by GroupNorm and ReLU activation. Spatial resolution is halved at each block via average pooling with a stride of 2. Rather than using only the final block as in the original architecture, the multi-scale readout extracts feature maps from every encoder block and passes them to the covariance pooling stage, so the final representation includes information from both early and deeper layers.

#### D. Covariance Pooling

Features are extracted independently from the output of each block. For each block, adaptive average pooling maps the feature tensor to a 2*×* 2*×* 2 spatial grid, producing *C×* 8 pooled features, where *C* is the number of channels. Simultaneously, a sample covariance matrix is computed using the first 32 channels, and only its upper-triangular elements are kept, providing 496 covariance features per block. Motivated by global second-order pooling in convolutional networks (24), this representation models the relationships among convolutional feature responses. In our setting, channel pairs that consistently co-activate across spatial locations produce high covariance values, allowing the representation to encode relational structure beyond average pooled activations. Block-wise features are concatenated to form a 9,664-dimensional feature embedding. The model contains 675,264 parameters and requires approximately 3 seconds per image on CPU.

#### E. Weight initialization

All network parameters were initialized using PyTorch’s default scheme (Kaiming uniform (25) for convolutional layers) with a fixed random seed of zero. Weights were frozen immediately after initialization.

## Experiments

We compared the predictive capabilities of the features extracted by the un-CNN to those extracted by state-of-theart structural MRI foundation models and by FreeSurfer, the traditional neuroimaging segmentation pipeline. All models were evaluated on sex classification, brain age regression, and BMI regression across three publicly available datasets (NKI, HBN, PPMI). NKI and HBN data were accessed through the Reproducible Brain Charts initiative (26), while PPMI data were obtained directly from the PPMI database (14). All datasets were previously collected and deidentified, and are available under the ethics approvals and data-use agreements of the originating studies. The present study was approved under CAMH Research Ethics Board protocols REB 2025/172, covering the younger samples, and REB 2025/174, covering older adults.

### A. Baseline Methods

#### Foundation Models

Four pre-trained brain MRI foundation models were evaluated for comparison. AnatCL (4) is a 3D ResNet-18 model trained using anatomy-guided weakly contrastive learning on 3,984 healthy subjects from OpenBHB (27). 3D-Neuro-SimCLR (5) is a self-supervised 3D ResNet-18 model trained with SimCLR on 44,958 T1-weighted scans from 11 public datasets. BrainIAC (6) is a Vision Transformer trained with SimCLR on 48,965 multiparametric MRI scans (T1, T2, and FLAIR) and ten clinical conditions across 34 datasets. Swin-Brain (7) is a SwinUNETR model trained using masked reconstruction and contrastive learning on 75,861 scans from private clinical data.

#### FreeSurfer

FreeSurfer (1) is a widely used neuroimaging pipeline for cortical surface reconstruction, anatomical parcellation, and morphometric quantification from T1-weighted MRI. Unlike TurboPrep, which produces spatially normalized volumes for downstream image-based feature extraction, FreeSurfer provides pre-computed regional anatomical measurements, such as cortical thickness and surface area, for predefined cortical regions. Cortical thickness and cortical surface area measurements derived from the Desikan–Killiany (aparc) (28) atlas (68 regions, 136 features) and the Schaefer (29) 400-parcel atlas (800 features) served as traditional baselines. For the NKI and HBN datasets, FreeSurfer-derived values were obtained from the Reproducible Brain Charts initiative (26) (FreeSurfer v6.0.1), whereas for PPMI, FreeSurfer v7.3.2 was used.

### B. Downstream Approach & Tasks

Each model produced one fixed-length feature vector per subject, which was evaluated using the same downstream pipeline. Model inputs differed according to each model’s requirements: the un-CNN and most foundation models used TurboPrep-processed volumes, AnatCL used CAT12 gray matter probability maps, and pre-computed morphometric features (cortical thickness and cortical surface area) for the FreeSurfer baselines. A Random Forest classifier or regressor served as the downstream model across all feature extractors and tasks, using *n*_estimators_ = 200, max_depth = 6, and min_samples_split = 5. For classification, max_features =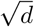 (where *d* denotes the feature dimensionality) and balanced class weights were used, whereas for regression, max_features = *d* was applied. Performance was assessed over five independent train/test partitions (90/10 split) using random seeds *{*0, 1, 2, 3, 42*}* . Results are reported as mean *±*standard deviation across 5 random seeds.

Three primary tasks were evaluated: sex classification (balanced accuracy), brain age regression (mean absolute error in years), and BMI regression (mean absolute error; NKI and HBN only).

### C. Datasets

Three publicly available datasets containing T1-weighted MRI images were used in this study. The Nathan Kline Institute Rockland Sample (NKI) dataset (12) included 958 subjects and represented a healthy lifespan cohort (357 male,601 female; age range 6–85 years, mean age 41.99 *±*20.22 years), with body mass index (BMI) available for 956 subjects (mean 26.5*±* 5.8). All NKI images were acquired at 1.0*×* 0.977*×* 0.977 mm resolution (176*×* 256 *×*256 voxels). The Healthy Brain Network (HBN) dataset (13) included 1000 subjects and represented a children/adolescent cohort enriched for learning and mental health disorders (519 male, 481 female; age range 5–22 years, mean age 11.3*±* 3.8 years), with BMI available for 962 subjects (mean 19.8*±* 5.1). HBN images were acquired with two protocols:1.0 mm isotropic (176 *×*256 *×*256) and 0.8 mm isotropic (224 *×*320 *×*320). In addition, the Parkinson’s Progression Markers Initiative (PPMI) dataset (14) included 894 subjects and represented an elderly clinical cohort (555 male, 339 female; age range 30–85 years, mean age 62.5 *±*10.0 years), comprising 711 individuals diagnosed with Parkinson’s disease and 183 healthy controls. PPMI images were acquired with heterogeneous protocols, resulting in 42 distinct matrix sizes and 24 distinct voxel resolutions ranging from 0.49 to 2.0 mm. All images were registered to a common MNI152 template space (193*×* 229 *×*193 at 1 mm isotropic) through the TurboPrep pipeline. 3D-Neuro-SimCLR and BrainIAC were not evaluated on the PPMI dataset due to the inclusion of PPMI data in their pretraining datasets.

All architectural and preprocessing choices for the un-CNN were optimized exclusively on the NKI dataset, using sex classification and brain age prediction as optimization targets. The HBN and PPMI datasets were used as held-out sets and BMI prediction was evaluated as an additional held-out task on NKI and HBN.

### D. Architecture Ablation

To quantify the contribution of each design choice to the performance of the un-CNN model, a cumulative ablation study was conducted on NKI dataset, which served as the optimization set for all architectural and preprocessing decision. Starting from the original un-CNN architecture (15), each modification was introduced sequentially and evaluated on sex classification and brain age regression: (1) multiscale readout from all blocks, (2) GroupNorm with average pooling, (3) percentile clipping with min-max normalization, (4) depthwise separable convolutions, (5) three-channel input construction, (6) double convolutions per block, (7) rank filtering, (8) tighter percentile clipping, (9) covariance pooling, and (10) robust intensity scaling.

## Results

### A. Un-CNN matched or outperformed baselines

Figure 2 summarizes the complete benchmark results across the three datasets and all evaluated tasks. Overall, the un-CNN matched or outperformed the evaluated foundation model and FreeSurfer baselines across datasets, despite using randomly initialized weights and no pretraining.

**Fig. 2.**
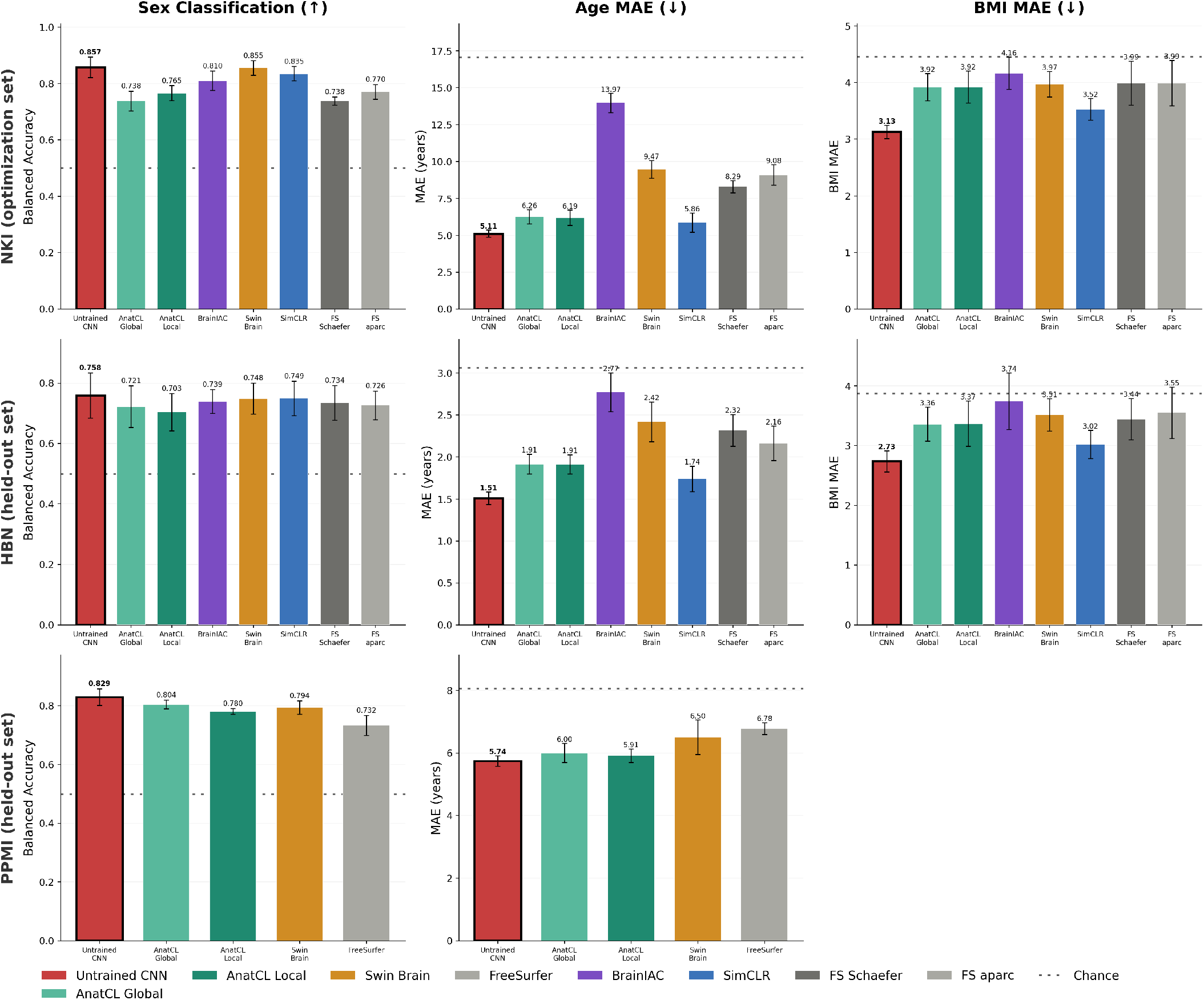
Untrained CNN (un-CNN) performance across datasets and downstream tasks.

On NKI, the un-CNN achieved the lowest age MAE (5.11 years) and BMI MAE (3.13), outperforming all foundation model and FreeSurfer baselines. For sex classification, it achieved 85.7% balanced accuracy, performing comparably to SwinBrain (85.5%) and exceeding 3D-Neuro-SimCLR (83.5%).

On HBN, the un-CNN achieved the best performance across all evaluated tasks: sex classification (75.8% balanced accuracy), age prediction (1.51 years MAE), and BMI prediction (2.73 MAE). Relative to the next-best model, 3D-Neuro-SimCLR, the un-CNN improved age prediction by 0.23 years and BMI prediction by 0.29 units. Importantly, HBN was not used during un-CNN architecture optimization, indicating good generalizability.

On PPMI, the un-CNN achieved the highest sex classification accuracy (82.9%) and the lowest age MAE (5.74 years), outperforming AnatCL (5.91 years) and FreeSurfer (6.78 years). PPMI was also entirely unseen during architecture optimization, further supporting the generalizability of the proposed architecture. As noted above, 3D-Neuro-SimCLR and BrainIAC were excluded from the PPMI evaluation because PPMI subjects were included in their pretraining datasets, illustrating a data leakage risk that untrained architectures avoid by design.

### B. Un-CNN substantially reduces computational costs

Table 1 compares deployment-time inference and reported training costs across different models. The final un-CNN achieved a total runtime of 2.68*±* 0.001 s/img, including timed preprocessing, feature extraction, and random forest prediction. Unlike foundation models, which require largescale pretraining before deployment, un-CNN uses fixed random weights and requires no training. In contrast, FreeSurfer requires several hours per subject for full `recon-all` reconstruction.

**Table 1.**
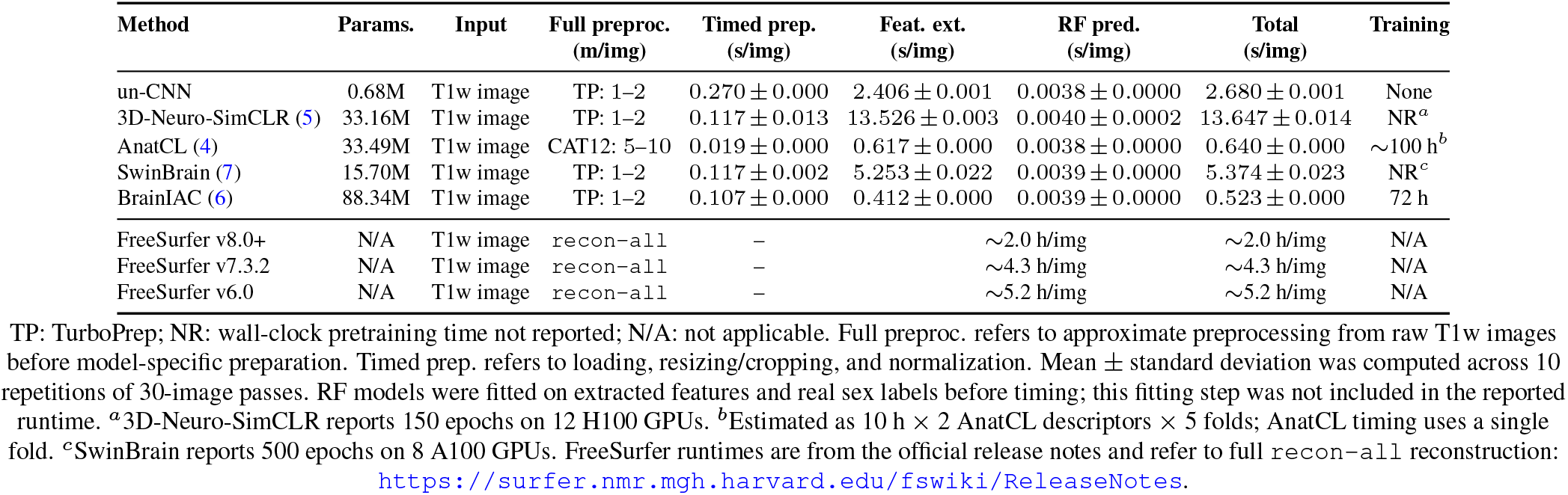
End-to-end computational-cost comparison on 30 NKI images across 10 repetitions. Runtime was measured using single-threaded DRAC CPU execution. RF fitting time is not included.

| Method | Params. | Input | Full preproc.<br>(m/img) | Timed prep.<br>(s/img) | Feat. ext.<br>(s/img) | RF pred.<br>(s/img) | Total<br>(s/img) | Training |
| --- | --- | --- | --- | --- | --- | --- | --- | --- |
| un-CNN | 0.68M | T1w image | TP: 1-2 | 0.270 ± 0.000 | 2.406 ± 0.001 | 0.0038 ± 0.0000 | 2.680 ± 0.001 | None |
| 3D-Neuro-SimCLR (5) | 33.16M | T1w image | TP: 1-2 | 0.117 ± 0.013 | 13.526 ± 0.003 | 0.0040 ± 0.0002 | 13.647 ± 0.014 | NR<sup>a</sup> |
| AnatCL (4) | 33.49M | T1w image | CAT12: 5-10 | 0.019 ± 0.000 | 0.617 ± 0.000 | 0.0038 ± 0.0000 | 0.640 ± 0.000 | ~100 h<sup>b</sup> |
| SwinBrain (7) | 15.70M | T1w image | TP: 1-2 | 0.117 ± 0.002 | 5.253 ± 0.022 | 0.0039 ± 0.0000 | 5.374 ± 0.023 | NR<sup>c</sup> |
| BrainIAC (6) | 88.34M | T1w image | TP: 1-2 | 0.107 ± 0.000 | 0.412 ± 0.000 | 0.0039 ± 0.0000 | 0.523 ± 0.000 | 72 h |
| FreeSurfer v8.0+ | N/A | T1w image | recon-all | — | ~2.0 h/img | ~2.0 h/img | ~2.0 h/img | N/A |
| FreeSurfer v7.3.2 | N/A | T1w image | recon-all | — | ~4.3 h/img | ~4.3 h/img | ~4.3 h/img | N/A |
| FreeSurfer v6.0 | N/A | T1w image | recon-all | — | ~5.2 h/img | ~5.2 h/img | ~5.2 h/img | N/A |

TP: TurboPrep; NR: wall-clock pretraining time not reported; N/A: not applicable. Full preproc. refers to approximate preprocessing from raw T1w images before model-specific preparation. Timed prep. refers to loading, resizing/cropping, and normalization. Mean ± standard deviation was computed across 10 repetitions of 30-image passes. RF models were fitted on extracted features and real sex labels before timing; this fitting step was not included in the reported runtime. <sup>a</sup>3D-Neuro-SimCLR reports 150 epochs on 12 H100 GPUs. <sup>b</sup>Estimated as 10 h × 2 AnatCL descriptors × 5 folds; AnatCL timing uses a single fold. <sup>c</sup>SwinBrain reports 500 epochs on 8 A100 GPUs. FreeSurfer runtimes are from the official release notes and refer to full recon-all reconstruction: https://surfer.nmr.mgh.harvard.edu/fswiki/ReleaseNotes.

Pretraining costs were obtained from the corresponding model papers when reported. For AnatCL, the reported time for a single training run was approximately 10 h; because the released evaluation includes two descriptors and five folds, we estimated the total pretraining cost as 10 h*×* 2*×* 5 = 100 h. BrainIAC directly reports a pretraining time of 72 h. For SwinBrain and 3D-Neuro-SimCLR, wall-clock pretraining time was not reported; therefore, we report only the available training configuration. SwinBrain was pretrained for 500 epochs using 8 NVIDIA A100 GPUs, whereas 3D-Neuro-SimCLR was pretrained for 150 epochs using 12 NVIDIA H100 GPUs with an effective batch size of 72. When available, costs are reported as wall-clock times rather than GPU-hours. Runtime benchmarking was performed on the Digital Research Alliance of Canada (DRAC) CPU infrastructure using single-threaded CPU execution, with all models evaluated on the same compute node. We set `OMP_NUM_THREADS=1`, `MKL_NUM_THREADS=1`, `OPENBLAS_NUM_THREADS=1`,and torch.set_num_threads(1). Runtime was mea-sured on 30 NKI images across 10 repetitions and includes model-specific timed preprocessing, feature extraction, and random forest prediction. Timed preprocessing includes input loading, resizing/cropping, and normalization. Random forest fitting time was not included.

### C. Ablation study identifies an efficient final un-CNN configuration

Table 2 and Figure 3 summarize the cumulative effect of each architectural and preprocessing modification on NKI. Overall, the ablation study shows that the optimized un-CNN sub-stantially improved age prediction relative to the default architecture while preserving strong sex-classification performance. Runtime was recorded for each cumulative variant as the total wall-clock time required to extract features for all NKI subjects and evaluate downstream sex classification and age regression across five random seeds. Thus, this runtime reflects the full model-selection/evaluation cost, including feature extraction and random forest fitting/evaluation, rather than deployment-time inference alone.

**Table 2.** Cumulative architecture ablation on NKI. Each row adds one modification to the previous configuration.

| Model | Final Dimension | Ablation runtime (s/img) | Sex Accuracy | Age MAE |
| --- | --- | --- | --- | --- |
| Default CNN(15) | 512 | 0.6 | 82.0% | 11.23 |
| +Multi-scale readout | 4,096 | 1.2 | 85.0% | 9.02 |
| +GroupNorm +AvgPool | 4,096 | 1.4 | 84.4% | 8.64 |
| +Clip [1,99] +MinMax | 4,096 | 1.3 | 86.1% | 7.72 |
| +Depthwise sep. conv | 4,096 | 4.5 | 85.1% | 7.19 |
| +3ch (unsharp+sobel) | 4,096 | 1.4 | 86.3% | 7.16 |
| +DoubleConv | 7,680 | 3.1 | 84.8% | 6.21 |
| +Rank filter | 7,680 | 2.8 | 85.1% | 6.19 |
| +Clip [2,98] | 7,680 | 2.8 | 85.5% | 6.17 |
| +Covariance pooling | 9,664 | 3.2 | 87.7% | 5.74 |
| **+Robust scaler** | **9,664** | **3.0** | **85.7%** | **5.11** |

**Fig. 3.**
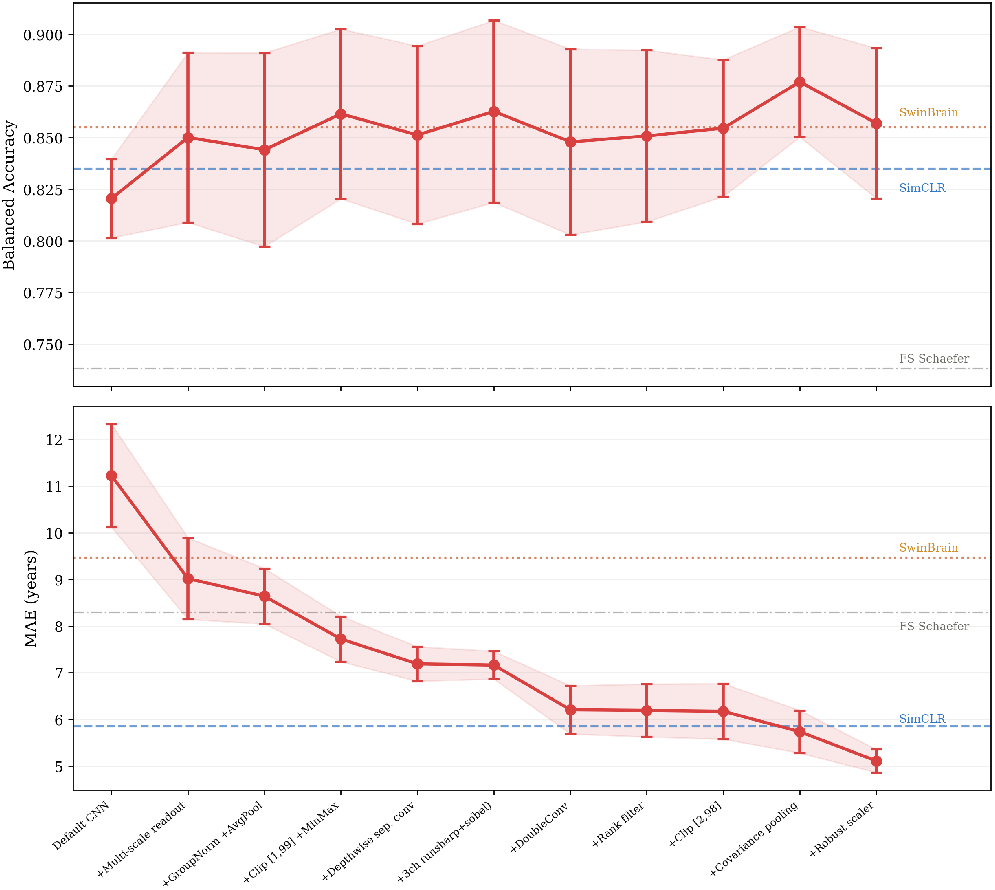
Cumulative architecture ablation on NKI (958 subjects). Top: sex classification balanced accuracy. Bottom: brain age prediction MAE (years). The trajectory shows the effect of each cumulative modification. Dashed lines indicate foundation model and FreeSurfer baselines evaluated on the same subjects.

#### Brain Age Prediction

The default untrained CNN (15) achieved an age MAE of 11.23 years. Multi-scale readout from all four convolutional blocks produced the largest early improvement, reducing MAE to 9.02 years, suggesting that information from intermediate spatial resolutions is important for subject-level MRI representations. Replacing BatchNorm with GroupNorm and MaxPool with average pooling, followed by percentile-based intensity clipping, further reduced MAE to 7.72 years. Sub-sequent architectural changes, including depthwise separable convolutions, three-channel preprocessing, and Double-Conv blocks, progressively improved age prediction, reducing MAE to 6.21 years and surpassing the FreeSurfer Schaefer baseline (8.29 years). Replacing unsharp masking with rank filtering and adjusting the clipping range to [2, 98] provided smaller additional gains.

The largest late-stage improvement came from covariance pooling, which reduced age MAE from 6.17 to 5.74 years. The final addition of robust intensity scaling further reduced MAE to 5.11 years, corresponding to a 54.5% relative improvement over the default architecture and a 12.8% improvement over 3D-Neuro-SimCLR (5.86 years).

#### Sex classification

showed a less monotonic pattern across the ablation sequence. The default CNN achieved 82.0% balanced accuracy, while several intermediate variants improved performance to the mid-80% range. Covariance pooling produced the highest sex classification accuracy (87.7%), whereas the final robust-scaling variant achieved 85.7%. Thus, while the final model was selected primarily for its strongest age-prediction performance, the ablation also shows that the proposed architectural changes maintained strong sex-classification performance throughout the optimization process.

## Discussion and Conclusion

Our results demonstrate that 3D untrained CNNs can match or exceed the predictive performance of state-of-the-art foundation models trained on thousands of brain MRI scans across multiple downstream tasks and datasets. Across all three datasets, the un-CNN architecture matched or exceeded the best foundation model on age prediction: 5.11 vs 5.86 years MAE on NKI (the optimization set), 1.51 vs 1.74 on HBN, and 5.74 vs 5.91 on PPMI — while requiring no data, no pretraining, and being fully reproducible from architecture code and a single random seed. These results generalized without modification from the NKI optimization sample to HBN and PPMI, spanning child and adolescent populations to elderly clinical populations.

Un-CNN consistently exceeded FreeSurfer-derived morphometric features across all tasks and datasets, while requiring only approximately 3 seconds per image on a single standard CPU, single thread compared to the hours of processing time needed by the FreeSurfer pipeline. This computational advantage is particularly relevant for large-scale neuroimaging studies and clinical settings where processing thousands of scans with traditional pipelines remains a practical bottleneck.

We attribute the effectiveness of un-CNN to three complementary factors. First, the CNN architecture itself imposes a strong inductive bias through local connectivity and hierarchical spatial compression, preserving meaningful anatomical structure even with random weights (9). Second, multichannel preprocessing (rank filtering and Sobel edge detection) compensates for the absence of learned first-layer filters by providing the network with hand-engineered texture and boundary information as input. Third, covariance pooling captures second-order spatial co-activation patterns between random feature maps (encoding which anatomical regions share similar local properties) providing substantially richer descriptors than average pooling alone. The cumulative ablation study (Table 2) confirms that each of these components contributes meaningfully, with covariance pooling and multiscale readout providing the largest individual improvements. Beyond predictive performance, un-CNNs benefit from the numerical stability properties of CNN inference. Gonzalez-Pepe et al. (30, 31) showed that CNN inference exhibits substantially lower numerical uncertainty than traditional neuroimaging processing pipelines such as FreeSurfer, while also identifying that training introduces additional variability through iterative stochastic optimization. Un-CNN eliminates this training-time variability entirely, by solely performing inference. This property is particularly valuable for reproducible neuroimaging research, where small numerical differences can propagate through analysis pipelines and affect downstream conclusions.

In summary, un-CNN is a fast, compute and storage efficient feature extractor that provides a practical alternative to both heavy foundation models and traditional morphometric pipelines for structural brain MRI analysis. The proposed architecture requires no training at all, yet matches or exceeds state-of-the-art performance across three datasets and multiple prediction tasks. Looking ahead, the computational efficiency of untrained CNNs makes them well-suited for largescale feature extraction across thousands of structural MRI scans in population-level neuroimaging studies. Extending the un-CNN architecture to functional MRI (fMRI) is also part of our future work.

## Notes

### Competing Interest Statement

The authors have declared no competing interest.

